# Astrocyte-Derived Exosomes Regulate Sperm miR-34c Levels to Mediate the Transgenerational Effects of Paternal Chronic Social Instability Stress

**DOI:** 10.1101/2024.06.17.599336

**Authors:** Alexandre Champroux, Mitra Sadat-Shirazi, Xuan Chen, Jonathan Hacker, Yongjie Yang, Larry A. Feig

## Abstract

The effects of chronically stressing male mice can be transmitted across generations by stress-specific changes in their sperm miRNA content that induce stress-specific phenotypes in their offspring. But how each stress paradigm alters the levels of distinct sets of sperm miRNAs is not understood. Here we describe evidence for **a**strocyte-derived **exos**omes (**A-Exos**) containing miR-34c mediating how chronic social instability (CSI) stress suppesses levels of miR-34c in sperm, which we showed previously contributes to how this stress protocol leads to both elevated anxiety and defective sociability in their female offspring and reduced sperm miR-34c in their male offspring. In particular, we found that CSI stress decreases content of miR-34c in A-Exos isolated from the prefrontal cortex and amygdala, as well as in blood of CSI-stressed males. Strikingly, miR-34c content is also reduced in A-Exos isolated from these tissues of their F1 male offspring, who also display reduced sperm miR-34c levels despite never being directly exposed to stress and transmit these stress related traits to their offspring. In addition, restoring A-Exos miR-34c content in the blood of CSI-stressed males by IV injection of miR-34c-containing A-Exos restores miR-34c levels in their sperm. These findings reveal a surprising role for A-Exos in maintaining sperm miR-34c levels by a process that when suppressed by CSI stress mediates this example of transgenerational epigenetic inheritance.

## 1 INTRODUCTION

The idea that traits acquired from environmental exposures can be transmitted to subsequent generations through epigenetic changes in their germ cells, a process referred to as transgenerational epigenetic inheritance [1], is well established in organisms like C. elegans, plants and fruitflies [2]. Evidence has been growing for this concept in rodents, including the effects of chemicals [3], drugs [4], diet [5], enriched environment[6], and chronic stress [7] where stress-induced changes in sperm miRNA content have been implicated. Human studies, based mainly on epidemiology are consistent with this concept [8], including summary statistics for eleven neuropsychiatric disorders that imply that the fraction of the heritability of these maladies not accounted for by classical genetics may be due to epigenetic inheritance [9].

A striking feature of these processes in mice that has yet to be explained is that each chronic stress paradigm studied alters the content of different sets of sperm miRNAs that lead to different behavioral effects in offspring. For example, Gapp et al [10] found that sperm of male mice exposed to unpredictable maternal separation at birth combined with unpredictable maternal stress (**MUS**) displayed elevated expression of a variety of sperm miRNAs, including miR-375, miR-200b, miR-672, and miR-466. And when all sperm RNAs from stressed or control mice were injected into control zygotes, male offspring from the former displayed elevated anxiety and depression, similar to what was observed in offspring of stressed males [10]. Rogers et al [11] showed that exposure of adult male mice to chronic variable stress (**CVS**) led to elevated levels of 8 different sperm miRNAs (plus miR-375 in common). And when all 9 were injected into control zygotes, it also recapitulated what was observed in offspring of stressed males. In this case no basal behavior changes were detected, but instead a blunted HPA axis response to stress was found in males and females.

In contrast, we demonstrated that exposure of adolescent male mice to chronic social instability (**CSI**) stress, which involves exposing mice to a novel mouse twice a week for 7 weeks generates elevated anxiety and defective sociability specifically in female offspring for at least three generations through the paternal lineage [12]. CSI stress leads to suppression in the sperm levels of miR-34/449 family members [13] as well as elevation of miR-409-3p [14], not only in directly stressed males but also in the sperm of their F1 male offspring who, despite never being exposed to stress, transmit the same stress associated traits to their male and female offspring. Moreover, suppressed sperm miR-34/449 levels in stressed mice are transmitted to **P**re**I**mplantation **E**mbryos (**PIEs**) upon fertilization [13]. We showed that this reduction in preimplantation embryo levels of miR-34/449 is critical for transmission of stress traits to offspring, since rescue of suppressed levels of these miRNAs reversed both the behavioral phenotypes in female offspring and reduced levels of miR-34/449 in sperm of male offspring [25].

Elevated stress hormones may play a role in how chronic stress promotes sperm miRNA changes, since injection of glucocorticoids can alter sperm miRNA levels. However, the miRNAs affected did not match any of those described above [15], implying a more specific signal likely participates. Moreover, others have detected changes in blood metabolites that appear to effect spermatogonia and mediate stress-induced metabolic effects in offspring, but their source or role in transmitting behavioral change to offspring was not shown [16]. Here, we describe experiments that show that at least for CSI stress, this more specific signal is in the form of reduced miR-34c in exosomes derived from astrocytes that appear to arise in a subset of brain regions known to respond to stress that are transmitted to through the blood to regulate its level in sperm.

## 2 METHODS

### 2.1 Animals and housing

All mice in this study were of the CD-1 strain obtained from Charles River Laboratories. Males used for CSI stress began the protocol at 28 days postnatal age, and control females used for breeding were 8 weeks of age. All animals were housed in temperature, humidity, and light-controlled (14hr on/10hr off LD cycle) rooms in a fully staffed dedicated animal core facility led by on-call veterinarians at all hours. Food and water were provided *ad libitum*. All procedures and protocols involving these mice were approved by the Institutional Animal Care and Use Committee of the Tufts University School of Medicine, Boston, MA.

### 2.2 Chronic social instability (CSI) stress

CSI protocol was conducted as described previously^6^. Briefly, the composition of each mouse cage (four mice per cage, 28 days old) was randomly shuffled twice per week for 7 weeks, such that each mouse was housed with three new mice in a fresh, clean cage. Control mice were housed with the same four mice per cage for the duration of the protocol. Any mice involved in physical interactions associated with this stressful condition are removed. After 7 weeks, mice were housed in pairs with a cage mate from the final cage change and left for 2 weeks to remove the acute effects of the final change. Mice were either sacrificed for sperm, blood, and brain collection or mated with random control female mice overnight to generate “F1” animals. We have previously shown that stressed males can be mated multiple times and still transmit stress phenotypes to their offspring [12], so male mice were used for mating or sperm collection as needed.

### 2.3 Mouse sperm collection

Mature, motile mouse sperm (from the cauda epididymis) were isolated via the swim-up method [17]. Briefly, male F0 and F1 mice were anesthetized under isoflurane and sacrificed via cervical dislocation. The caudal epididymis and vas deferens were dissected bilaterally and placed in 1mL of warm (37°C) M16 medium (Sigma-Aldrich, M7292) in a small Petri dish. Under a dissection microscope, sperm were manually expressed from the vas deferens using fine forceps, and the epididymis was cut several times before incubating at 37°C for 15 min to allow mature sperm to swim out, then large pieces of tissue were removed. The remainder of the extraction took place in a 37°C warm room. The sperm-containing media was centrifuged at 3,000 RPM for 8min, the supernatant was withdrawn and discarded, and 400μL of fresh, warm M16 medium was carefully placed on top of the pellet. The tubes were then allowed to rest at a 45° angle for 45 min to allow the motile sperm to swim-up out of the pellet into the fresh medium. The supernatant containing the mature sperm was carefully withdrawn and centrifuged again for 5 min at 3,000 RPM to pellet the motile sperm. The supernatant was withdrawn and discarded, and the pellet was frozen on dry ice for later processing.

Caput sperm were collected as described[18].Two small incisions were made at the proximal end of caput and using a 26G needle holes were made in the rest of the tissue to let the caput epididymal fluid ooze out. Sperm-containing media was incubated for 15 minutes at 37°C, then transferred to a fresh tube and the sperm-free epididymis tissues were directly frozen in liquid nitrogen and stored at −80°C. The sperm were incubated for another 15 minutes at 37°C, then sperm were collected by centrifugation at 2,000 x g for 2 minutes, followed by a 1X PBS wash, and a second wash with somatic cell lysis buffer (low SLB buffer, 0.01% SDS, 0.005% Triton X-100 in PBS) for 10 minutes on ice to eliminate somatic cell contamination. Somatic cell lysis buffer treated sperm were collected by centrifugation at 3,000xg for 5 minutes, and finally washed with 1X PBS before freezing down.

### 2.4 Epididymosome isolation by ultracentrifugation and density-gradient

After pelleting cauda sperm following the swim-up protocol in M16 medium (Sigma, M7167), the supernatant was centrifuged at 2,000xg for 10min, 10,000xg for 30min and then ultracentrifuged at 120,000xg at 4°C for 2h. The epididymosomal pellet was then washed in PBS 1X at 4°C and ultracentrifuged at 120,000xg at 4°C for 2h. The resulting pellet was resuspended in 50µl of PBS [19].

### 2.5 Submandibular blood collection

The submandibular blood was collected with K2E (Bd microtainer) from the submandibular vein using an animal lancet (size 5.5 mm, Goldenrod, Medipoint). Then the blood was centrifuged at 2,000xg, for 15min at 4°C before frozen for RNA extraction.

### 2.6 Brain dissection and exosome extraction from brain

Animals were anesthetized with isoflurane, then the brain was then extracted and immediately placed in a solution of ice-cold sterile phosphate-buffered saline (PBS). The prefrontal cortex (PFC), hippocampus, amygdala, and somatosensory cortex were then dissected according to the Paxinos atlas [20]. For exosome extraction, each brain region from four animals within the same experimental group was combined to ensure a representative pool. Exosomes were isolated from brain tissue as described [21]. Tissue was first digested with an enzyme solution containing 2 mg/ml collagenase in Hibernate E for 15 minutes at a temperature of 37°C. After digestion, a protease/phosphatase inhibitor was added in a volume equal to two times that of the total sample volume. The mixture underwent sequential centrifugations at specific speeds (300g for 10 minutes, 2,000g for 10 minutes, and 10,000g for 25 minutes) at 4°C. The desired exosome-containing supernatant was carefully collected. To further refine the exosome isolation, the collected supernatant was passed through a 0.45 μm filter. The exosomes were subsequently extracted using qEVorigincal 35mm Gen2 columns (Izon Sciences). Specific fractions (7-9) from the column elution were collected, concentrating the exosome samples using a 100 kDa cut-off tube. The correct vesicle diameter size for exosomes (∼100nm) was confirmed by Zetaview particle analysis see **Suppl Fig. 1**.

### 2.7 Isolation of exosomes from plasma

Blood was collected by cardiac puncture of the heart of the mice before sacrifice. Then the blood was centrifuged at 2,000xg for 15min at 4°C before frozen. Total exosomes from the plasma were purified using the Total exosomes isolation kit (Invitrogen, #4484450). Briefly, plasma was thawed at 36°C and then centrifuged at 2,000 x g for 20 minutes, followed by 10,000xg for 20 minutes. Then, 0.5 volume of 1X PBS was added followed by 0.05 volume of proteinase K and the sample was incubated at 37°C for 10 minutes. The exosomes precipitation reagent was then added at 0.2 volume, incubate at 4°C for 30 minutes and centrifuged at 10,000 x g for 30 minutes at 4°C. The correct vesicle diameter size for exosomes (∼100nm) was confirmed by Zetaview particle analysis see **Suppl Fig. 1**.

### 2.8 Isolation of Astrocyte Exosome by Immunoprecipitation

Astrocyte-derived exosomes (A-Exo) were purified from total exosome fractions from the brain and blood as described [22]. Briefly, samples were incubated with 3µg anti-GLAST (ACSA-1) biotinylated antibody (Miltenyi Biotec, USA) and BSA (3%) for 4 hours at 4°C after which streptavidin-agarose UltraLink Resin (Thermo Fisher Scientific) for 1 hour at 4°C. After a brief centrifugation (200g, 10 min, 4°C), the supernatant and pellet were used for RNA extraction.

### 2.9 Primary astrocyte culture and exosome purification

Our published procedure was used[23] Briefly, primary astrocyte cultures, were derived from P0-P3 mouse brain. Exosomes were prepared from 72hrs astrocyte conditioned medium (ACM) that was serum free. ACM was first spun at 300 × g for 10 min at room temperature to remove suspension cells, then at 2000 × g for 10 min at 4 °C to remove cell debris, then underwent following purification steps or stored at −80 °C. ACM supernatant was first concentrated (to 500 μl) by centrifugation at 3500 × g for 30 min at 4 °C using Centricon® Plus-70 Centrifugal Filter Devices with a 30k molecular weight cutoff (MilliporeSigma). Then the concentrated supernatant was passed through a 0.22 μm PES filter. The qEVoriginal 35 nm columns Gen2 (Izon Science, MA, USA) were then used as described above. Finally, the purity of exosomes was confirmed by tunable resistive pulse sensing.

### 2.10 Astrocyte-derived exosome injection

Astrocyte-derived exosomes (10ug protein) or saline controls were injected in the tail vein at day D0, D3 and D5 in a final volume of 100µl.

### 2.11 RNA extraction, cDNA synthesis and Real-time PCR

Total RNA, including miRNA, was extracted from A-Exos and the remaining exosomes using single-cell RNA extraction kit (Norgen Biotek, Canada), according to manufacturer protocol. Concentration of RNA was determined using a NanoDrop-1000 (Thermofisher Scientific). 10 ng of RNA from each sample was reverse transcribed using TaqMan Advanced miRNA cDNA Synthesis Kit (ABI, USA). Real-time PCR was performed for each target and sample in duplicate on a StepOnePlus PCR System (Applied Biosystems) with primers from TaqMan™ Advanced miRNA Assay. All data were analyzed using the Comparative ΔΔCT method to generate relative expression data using an endogenous miRNA that does not change under conditions used in these experiments as the internal control for miRNAs, as is standard procedure [24]. Supplementary Fig. 3 confirms that miR-192 is a suitable internal control for these experiments as it was for our previous study on CSI stress and miR-34c [25].

### 2.12 Protein Extraction and Western Blotting

Protein extraction from exosomes was carried out using RIPA lysis buffer containing 50 mM Tris-base at pH 7.5, 0.1% sodium dodecyl sulfate, 0.5% sodium deoxycholate, and 150 mM sodium chloride. The protein concentration was determined using NanoDrop spectrophotometry. Electrophoresis was carried out on a 10% polyacrylamide gel, and the separated proteins were transferred onto a polyvinylidene fluoride membrane (Bio-Rad, USA). Overnight incubation at 4°C with a primary antibody (GLAST1, CD81, TSG101 and CD63ABclonal, USA) at a 1/1000 dilution in skim milk followed by a one-hour incubation with a secondary antibody (Goat anti-Rabbit, ABclonal, USA, 1/3000) at room temperature. The blots were then washed three times with TBST, and visualization of protein bands was achieved using enhanced chemiluminescence (ECL, Bio-Rad, USA) in Chemidoc (Bio-Rad, USA).

### 2.13 Experimental Design and Statistical analysis

GraphPad Prism 9.0 software package was used for analyzing qPCR data. For data comparing only two groups, Mann-Whitney test was used. p ≤ 0.05 was considered significant.

## 3 RESULTS

### 3.1 Cauda sperm and epididymosome and levels of miR-34c are suppressed by CSI stress

A variety of sperm non-coding RNAs, including miRNAs, have been shown to be added to sperm via RNA-carrying exosomes secreted by epithelial cells of the cauda (final) section of the epididymis, called epididymosomes. And the RNA content of sperm has been shown to be altered when the environment changes epididymosome RNA content [26]. To begin to understand how CSI stress controls sperm miRNA levels we investigated whether this pathway is relevant to its effects. **Fig. 1** shows that it is, since miR-34c levels are not altered by CSI stress in sperm that just left the testes and entered the caput epididymis **(Fig.1A)** but are reduced 5-6-fold in sperm (p=0.0365) **(Fig. 1B)** and epididymosomes **(Fig. 1C)** purified from the cauda epididymis (p=0.0489).

**Figure 1:**
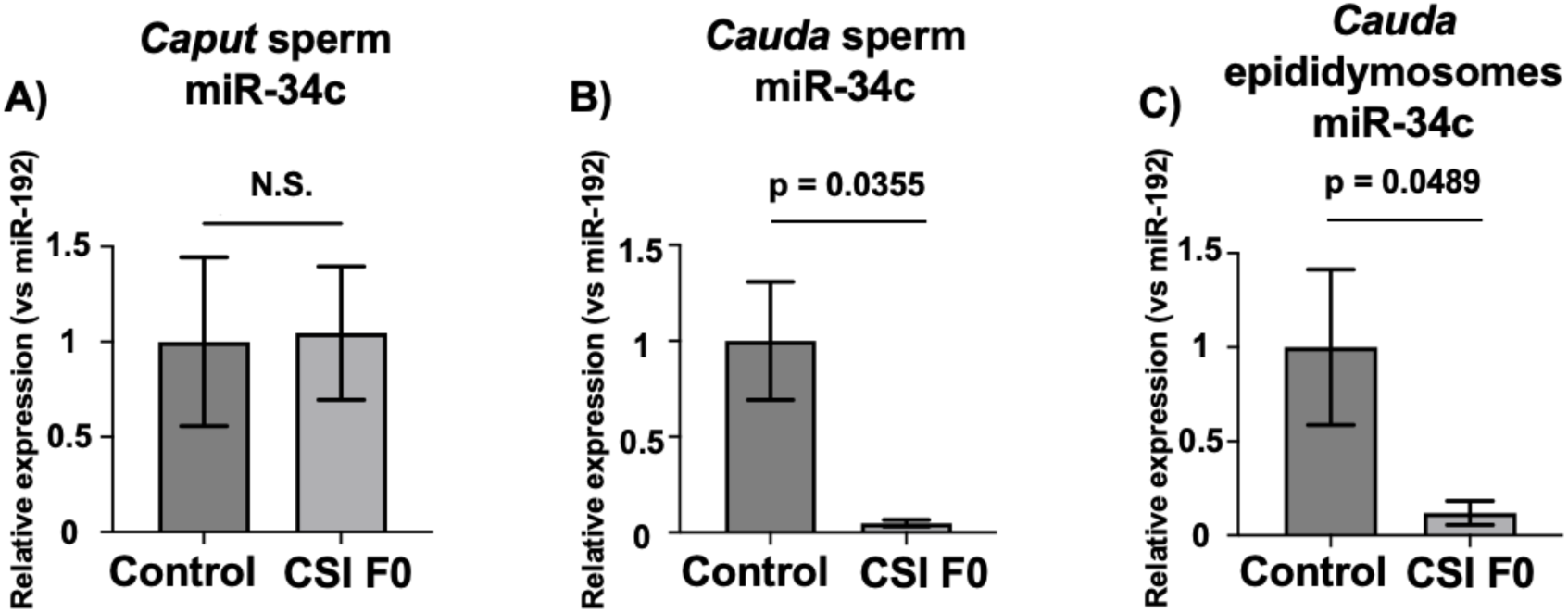
CSI stress reduces miR-34c levels in epididymosomes and sperm isolated from the *cauda* epididymis, but not sperm isolated from the *caput* epididymis. miR-34c levels in sperm isolated from *caput* (A) and *cauda* (B) regions of the epididymis from control and CSI stressed males. **C)** miR-34c levels in epididymosomes isolated from the *cauda* epididymis of control and CSI stressed males. n=5 for *caput* and *cauda* sperm and n=4 for *cauda* epididymosomes. Data are expressed as mean ± S.E.M; Mann-Whitney test, Two-tailed, WT: wild-type, CSI: Chronic Social Instability. Each point represents an individual mouse. All values are compared to the internal control miR-192 that does not change enough by any perturbation to significantly alter the conclusions (see **Suppl. Fig. 3**) and control ratios are set to 1.

### 3.2 CSI stress suppresses blood astrocyte-derived exosome content of miR-34c across generations

The idea that there may be a specific signal from the brain to germ cells that contributes to the transmission of acquired traits to offspring is supported by studies in *c. elegans* that show neurons send a RNA signal to germ cells to carry out this function[27]. That it could be derived from astrocytes was suggested by the fact that astrocytes are highly responsive to stress and the proof of principle study in mammals showing that when a human miRNA was expressed specifically in brain neurons and astrocytes of mice by viral infection, the human miRNA was subsequently found in the epididymis of males and offspring after their mating [28]. The possibility that the stress sensitive signal could be miR-34c itself was suggested by the fact that miRNAs are known to be carried in blood by exosomes and miR-34c has already been detected in them [29].

We began by testing **a**strocyte-derived **exos**omes (**A-Exos**) for this function using a well-characterized protocol to purify them from the total pool of isolated blood exosomes by immunoprecipitation using antibodies to the astrocyte-specific surface marker Glast1 [22]. **Fig. 2A** shows that this system works as expected as immunoprecipitation of the total pool of exosomes with Glast1 antibodies co-immunoprecipitated the exosome marker CD81 and omitting the Glast1 antibody led to undetectable levels of CD81 on the beads. miR-34c was also undetectable (data not shown). **Fig. 2B** reveals that miR-34c levels in A-Exos purified from the blood of CSI-stressed males were ∼5-fold lower (p=0.0389) than that found in A-Exos from control mice. Importantly, a similar, if not greater, decline in miR-34c A-Exos content (p=0.0055) was also found in the F1 male offspring of CSI stressed males that also displayed reduced miR-34c in their sperm. In contrast, miR-375, whose level rises in sperm after exposure of males to different types of chronic stress that produced different effects in offspring [10],[11] did not change across generations (**Fig. 2C**). Interestingly, no significant change was observed in blood A-Exos miR-34c content in F1 female offspring of CSI stressed males (see **Suppl Fig. 2**) consistent with our original observation that transmission through the female lineage faded across generations suggesting its transmission was not through the germline[12].

**Figure 2:**
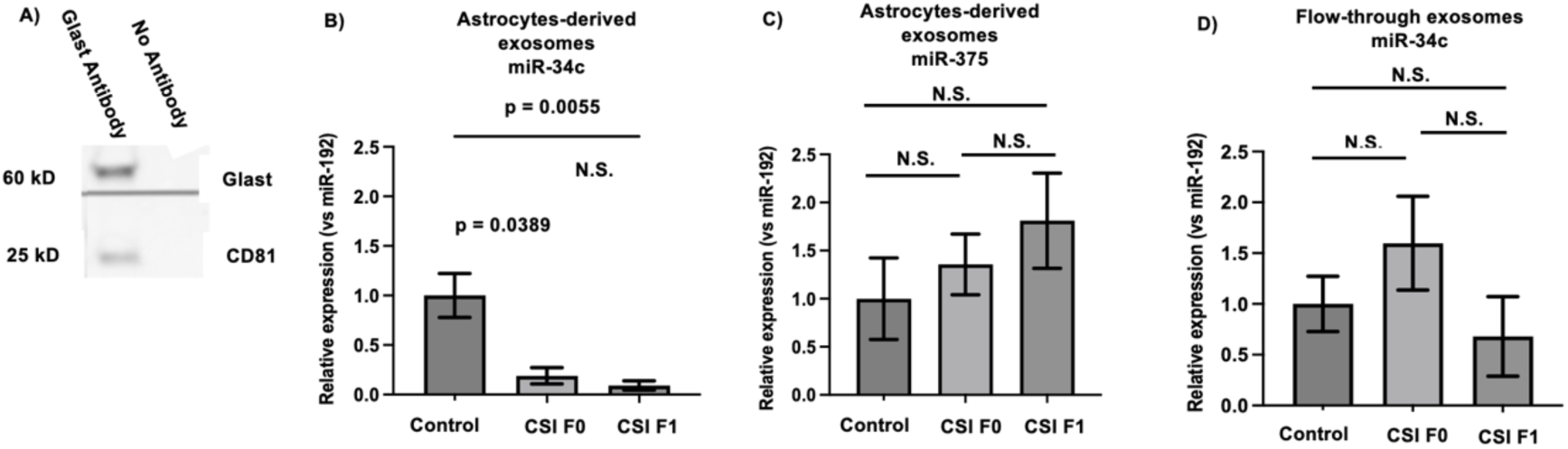
CSI stress suppresses the content of miR-34c in male, blood astrocyte-derived exosomes across generations. **A)** Co-IP of CD81 on western blot after immunoprecipitation with either anti-Glast antibody (Ab) or without (No Ab). **B)** Levels of miR-34c in astrocytes-derived exosomes immunoprecipitated from total plasma exosomes from CSI stress F0 males and their F1 male offspring. F(2,19) =7.878, p=0.003, Anova. N=7 for control and CSI F0 and N=8 for CSI F1 **C)** Levels of miR-375 in astrocytes-derived exosomes isolated from plasma of CSI stressed males and their F1 male offspring. F(2,19) = 0.1771, N=6 for control and N=7 for CSI F0 and F1; **D)** Levels of miR-34c in the non-precipitated flow-through after immunoprecipitation of astrocytes-derived exosomes. F(2,17)= 0.3063, Anova. N=6 for control CSI F0 and N=8 for CSI F1. All values are compared to the internal control miR-192 that does not change enough by any perturbation to significantly alter the conclusions (see **Suppl. Fig. 3**) and control ratios are set to 1.

miR-34c was also detectable in the non-precipitated fraction of blood exosomes that contains exosomes from other cell types in the brain and peripheral tissues. However, its level was not affected by CSI stress (**Fig. 2D**) supporting the concept that it is predominantly A-Exos that mediate a miR-34c signal between the brain and sperm through the blood. This suggest that CSI stress likely decreases the content of miR-34c in a similar number of secreted A-Exos, an idea also supported by finding similar levels of miR-375 in control and stressed fractions. However, if only a small fraction of A-Exos in the blood carry significant amounts of stress-responsive miR-34c, reduced levels of that fraction of exosomes could be masked.

In contrast, miR-449a, which we found is also suppressed in sperm of CSI stressed mice and in sperm of their male F1 offspring was not detectable in blood A-Exos (data not shown) suggesting its levels in sperm may be regulated differently. However, we showed recently that reduced sperm miR-34c levels are sufficient for transmitting traits to offspring, because when we transiently rescued reduced miR-34c in preimplantation embryos derived from CSI stresses males it also restored miR-449a levels by increasing its expression from the miR-449 early embryo gene [25]. Thus, if the same process occurs in the epididymis, suppression of miR-34c containing A-Exos might also lead to reduced levels of miR-449 in epididymosomes and then sperm, a topic presently under investigation.

### 3.3 Reduced levels of A-Exos miR-34c levels in blood of CSI stressed males are responsible for the reduced levels of miR-34c in sperm

To show that CSI-induced suppression of miR-34c levels in blood A-Exos is a necessary step in how it controls sperm miR-34c levels, we elevated miR-34c levels in blood exosomes of stressed males by IV injection of miR-34c-containing A-Exos purified from cultured mouse astrocytes (as described in [30]) and tested whether this restored sperm miR-34c levels. The feasibility of testing the biological activity of miR-34c in A-Exos by IV injection was supported by previous experiments showing that it could protect against brain injury induced by cerebral ischemia/reperfusion injury [29].

Mouse sperm reside for as much a week in the cauda epididymis where they absorb epididymosomes. Thus, we did a set of pilot experiments to determine how many A-Exos and how many injections we needed to restore blood miR-34c levels in CSI stressed mice to normal and settled on 1ug exosome protein on day 0, 3 and 5 (**Fig. 3A)**. Then, we repeated the experiment injecting either saline or A-Exos as described and measured the average levels of blood miR-34c on day 4 and 7 (**Fig. 3B**), and epididymosome and sperm miR-34c levels on day 7. **Fig. 3B-D** shows that when we restored blood miR-34c levels we also restored it in cauda epididymosomes and sperm, demonstrating that the level of miR-34c in blood A-Exos does control its level in sperm.

**Figure 3:**
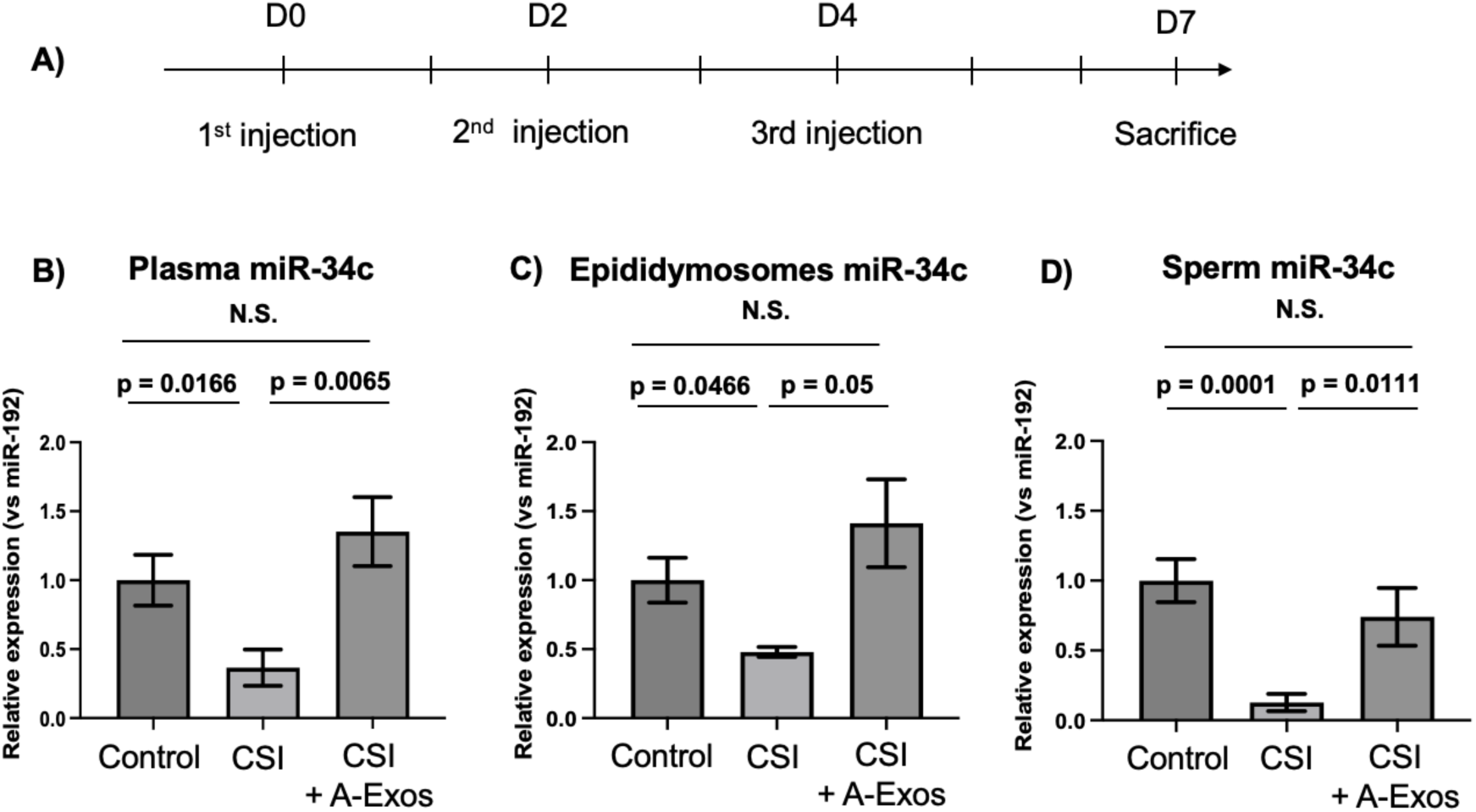
Elevation of blood levels of A-Exos miR-34c in CSI stressed males restores sperm levels of miR-34c. **A)** Scheme for injection of astrocytes-derived exosomes or saline into CSI stressed males and blood collection. **B)** Plasma levels of miR-34c of unstressed (control) males, and the average plasma levels of miR-34c sampled at day 4 and 7 from saline injected (CSI) or exosome injected (CSI+A-Exos) CSI stressed mice; N=7 for unstressed and N=4 for CSI stressed mice. **C)** Levels of miR-34c in epididymosomes isolated from unstressed males (control), and CSI stressed males injected with either saline (CSI) or purified astrocyte-derived exosomes (CSI + A-Exos) Welch’s T-test was used. N= 4 for control, saline and exosome injected mice. **D)** The same as C except miR-34c levels were assayed in cauda sperm; Welch’s T-test was used. N= 12 for control and N=4 for saline or exosome injected mice CSI stressed mice. Data are expressed as mean ± S.E.M; CSI: Chronic Social Instability. Each point represents an individual mouse. All values are compared to the internal control miR-192 that does not change enough by any perturbation to significantly alter the conclusions (see **Suppl. Fig. 3**) and control ratios are set to 1.

### 3.4 CSI stress reduces level of A-Exos miR-34c in the pre-frontal cortex and amygdala, but not hippocampus, of both stressed males and their F1 male offspring

To reveal where in the brain the depleted miR-34c-bearing A-Exos in the blood of CSI stressed males and their F1 male offspring might derive, we assayed miR-34c levels in A-Exos purified from three brain regions known to respond to chronic stress, and one that is not. We first isolated the total exosome fraction from the adult brain as described [21] and then used Glast1 antibodies to immune-purify the A-Exos fraction. And just as in studies of blood A-Exos in Fig. 2, immunoprecipitation of A-Exos from total pools of exosomes was specific because miR-34c levels were undetectable when immunoprecipitations were done without Glast1 antibodies. We began with the **p**re**f**rontal **c**ortex **(PFC)** because CSI stress has already been shown to alter its transcription profile [31]. **Fig. 4A** demonstrates ∼20-fold decline in the content of miR-34c in A-Exos isolated from both directly stressed mice (p=0.02) and their F1 male offspring (p=0.009). And like A-Exos from the blood, no change was detected in either A-Exos population for miR-375, whose level in sperm is elevated by different stress paradigms that produces different effects in offspring [10], [11]. As in blood, miR-34c was detectable in exosomes in the flow-through non-precipitated fraction from these other sources. But its level was not changed by CSI stress exposure in neither directly stressed mice nor their F1 male offspring (**Fig. 4B**), consistent with the model that miR-34c in A-Exos are the major regulators of stress induced changes in sperm miR-34c content. Also, like A-Exos in blood, miR-449a was not detectable in A-Exos isolated from the PFC (data not shown).

**Figure 4:**
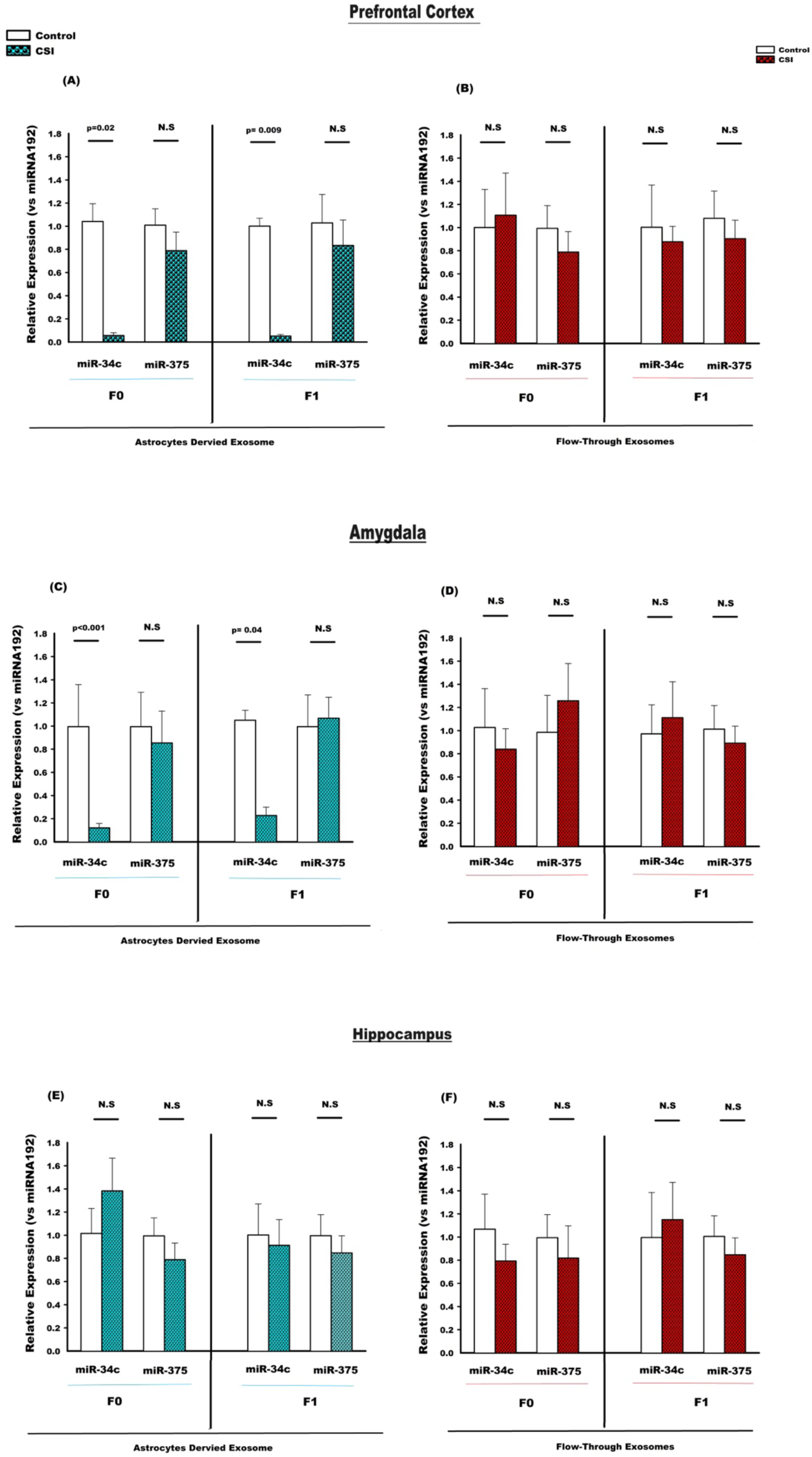
CSI suppresses levels of miR-34c in A-Exos isolated from the prefrontal cortex and amygdala, but not hippocampus, across generations. **A-B) Prefrontal Cortex-A)-Left-**miR-34c and miR-375 levels in A-Exos immune-purified from the total pool of exosomes in the prefrontal cortex of control and CSI-stressed F0 males (n=7), **Right-**the same analysis on A-Exos from their F1 male offspring (n=9 for miR-34c, n=5 for miR-375). **B) Left**-miR-34c and miR-375 levels in the non-precipitated, flow-through fraction of exosomes purified from the prefrontal cortex of control and CSI-stressed F0 males (n=5), Right-the same analysis on exosomes from their F1 male offspring (n=5) **C-D**) **Amygdala-C)-Left-** miR-34c and miR-375 levels in A-Exos immune-purified from the total pool of exosomes in the amygdala of control and CSI stressed males (n=6) **Right-**the same analysis on A-Exos from their F1 male offspring (n=7); **D) Left**-miR-34c and miR-375 levels in the non-precipitated, flow-through fraction of exosomes purified from the amygdala and CSI-stressed F0 males (n=5), Right-the same analysis on exosomes from their F1 male offspring (n=5). **E-F) Hippocampus**- **E) Left-**miR-34c and miR-375 levels in A-Exos immune-purified from the total pool of exosomes in hippocampus of control and CSI-stressed F0 males (n=5), **Right-**the same analysis on A-Exos from their F1 male offspring (n=5). **F) Left**-miR-34c and miR-375 levels in the non-precipitated, flow-through fraction of exosomes purified from the hippocampus of control and CSI-stressed F0 males (n=5), **Right**- the same analysis on exosomes from their F1 male offspring (n=5). Bars represent the fold differences in mean normalized expression values, and the error bars are standard errors (+/- SEM). Each data point is a pool of tissues from 4 mice. All values are compared to the internal control miR-192 that does not change enough by any perturbation to significantly alter the conclusions and control ratios are set to 1.

miR-34c levels in A-Exos isolated from the amygdala were also suppressed by CSI stress, although to a lesser degree (∼5-fold), across generations of males (**Fig. 4C** p<0.001 for F0, and p=0.04 for F1). As in the PFC, the non-astrocyte derived exosome fraction expressed significant miR-34c, but its level was also not affected by exposure of mice to CSI stress (**Fig. 4D**).

In contrast to the PFC and amygdala, the level of miR-34c in A-Exos isolated from the hippocampus was not suppressed by exposure of males to CSI-stress, nor in their male offspring (**Fig. 4E**). Like the PFC and amygdala, the non-astrocyte-derived exosome fraction also contained significant levels of miR-34c, and its level was not altered by CSI-stress (**Fig. 4F**), indicating that a different exosome population in the hippocampus is not involved in the CSI response. Finally, we found that A-Exos isolated from the sensory cortex, not known to be stress-responsive, did not even contain detectable levels of miR-34c (data not shown).

## 4 DISCUSSION

There is growing possibility that a significant fraction of inherited susceptibility to mental health disorders derives from one’s parent’s experiences transmitted to them via epigenetic changes in parent’s germ cells [32]. Much of this evidence derives from mouse models of chronic stress where stress-specific changes in sperm miRNA content leads to stress-specific alterations in offspring phenotypes. A mechanism underlying how a specific stressor alters the content of a specific sperm miRNA has yet to be revealed, until now at least for CSI stress. The experiments described here show that CSI stress reduces sperm levels of miR-34c by suppressing the miR-34c content carried by A-Exos in blood, since we demonstrated that restoring blood A-Exos miR-34c content in CSI-stressed mice restored miR-34c levels in their sperm. We implicated the prefrontal cortex and amygdala as at least some of the sources of miR-34c-containing A-Exos that are suppressed by CSI stress since the level of A-Exos miR-34c in these brain regions was severely reduced not only in CSI-stressed males but also in their F1 male offspring, both of which display reduced levels of miR-34c in their blood A-Exos and sperm. But until we can define the specific astrocyte populations in these brain regions that are involved and test the effect of manipulating their exosome function directly, we must depend upon this strong correlative evidence. Interestingly, another stress-responsive brain region, the hippocampus, contains significant levels of A-Exos miR-34c but its levels are not affected by CSI stress. Presumably these exosomes may be used for a more local function of miR-34c or respond to a different stressor not yet reported that also functions through miR-34c in A-Exos to affect the next generation.

These experiments also revealed the surprising finding that A-Exos are normally responsible for sustaining the level of miR-34c in sperm. And when CSI stress inhibits miR-34c levels carried by them in the blood, presumably by suppressing their output from astrocytes in the prefrontal cortex and amygdala (and possibly other brain regions), suppressed sperm miR-34c levels are the result. Moreover, these A-Exos in blood may regulate miR-34c levels in other stress-responsive tissues, and thus be part of a novel mechanism to alter peripheral tissue function in a stress-specific manner.

What remains to be determined besides identifying the specific astrocyte populations involved, is how CSI stress alters their function in the PFC and amygdala, but not hippocampus, such that either less miR-34c is uploaded into each exosome produced or less miR-34c-containing exosomes are generated. For the former case, CSI stress may suppress cellular miR-34c uploading into existing exosomes by either suppressing cellular miR-34c levels or by inhibiting regulators that upload specific miRNAs into exosomes (for review see [33]). Alternatively, the seven weeks of CSI stress beginning during adolescence may lead to the loss of the specific astrocyte populations in the PFC and amygdala that are sensitive to CSI stress to explain the large ∼20-fold decline in miR-34c levels. Why this may occur in the PFC but not hippocampus could be due to the difference in their susceptibility to chronic stress during adolescence. In particular, numerous studies [34–37] have demonstrated that stress induces dendritic retractions and synaptic losses in both adolescent and adult medial PFC. Notably, dendritic branching can recover to pre-stress levels when adult subjects are removed from stressful conditions [38]. However, such recoveries are not observed following adolescent social instability stress, suggesting that adverse experiences during adolescence, when the CSI stress we impose begins, may permanently disrupt typical PFC astrocyte development [31]. Interestingly this effect was not observed in the hippocampus [39]. Moreover, there is a significant enhancement in amygdala-PFC connectivity during adolescence [40]. Thus, the changes in A-Exo miR-34c cargo that we observed only in the PFC and amygdala might be attributed to the fact that this connectivity undergoes late maturation, making it more vulnerable compared to other regions like the hippocampus.

Another striking finding is that astrocytes in the same brain regions in F1 offspring of stressed mice “inherit” the same defect in exosome function their fathers acquired by exposure to CSI stress. This can explain why we also observe reduced A-Exos miR-34c levels in their blood and sperm, so that they can transmit the same stress related traits to their offspring. These astrocytes were never directly exposed to stress, but likely acquired this phenotype because of improper astrocyte development that arose due to reduced miR-34c during their preimplantation embryo stage, a defect we showed contributes to reduced miR-34/449 levels in their sperm [41].

We also reported the involvement of another sperm miRNA in this example of epigenetic inheritance, miR-409-3p, whose level in sperm rises in mice exposed to CSI stress as well as in their male offspring [14]. However, we could not detect it in A-Exos from the PFC or blood (data not shown). This raises the possibility that miRNAs whose levels rise in response to chronic stress are regulated by different mechanisms. This concept is supported by our finding that miR-375, whose sperm levels rise in response to both unpredictable maternal separation at birth combined with unpredictable maternal stress and chronic variable stress[10, 11], is present in A-Exos from the PFC and amygdala, but unlike miR-34c its level does not change in response to CSI stress. Interestingly, we also proposed that miRNAs whose sperm levels rise in response to chronic stress function differently at another step in the epigenetic inheritance process; how stress-induced change in their very low content in sperm can have a large impact on early-embryos upon fertilization [41].

These findings raise several related implications of interest to pursue in the future. For example, there is growing interest in using brain-derived blood exosome content as a biomarker for various neurological disorders [42]. The results presented here raise the possibility that reduced A-Exos miR-34c content in blood A-Exos could be an indicator of early life trauma. This possibility is supported by our previous study indicating that miR-34c is also reduced in sperm of men who were exposed to a high level of Adverse Childhood Experiences (ACEs) [13], a finding subsequently confirmed using a different scale of early life trauma [43].

## Funding

NICHD

## Supplementary Figure Legends

**Supplemental figure 1:**
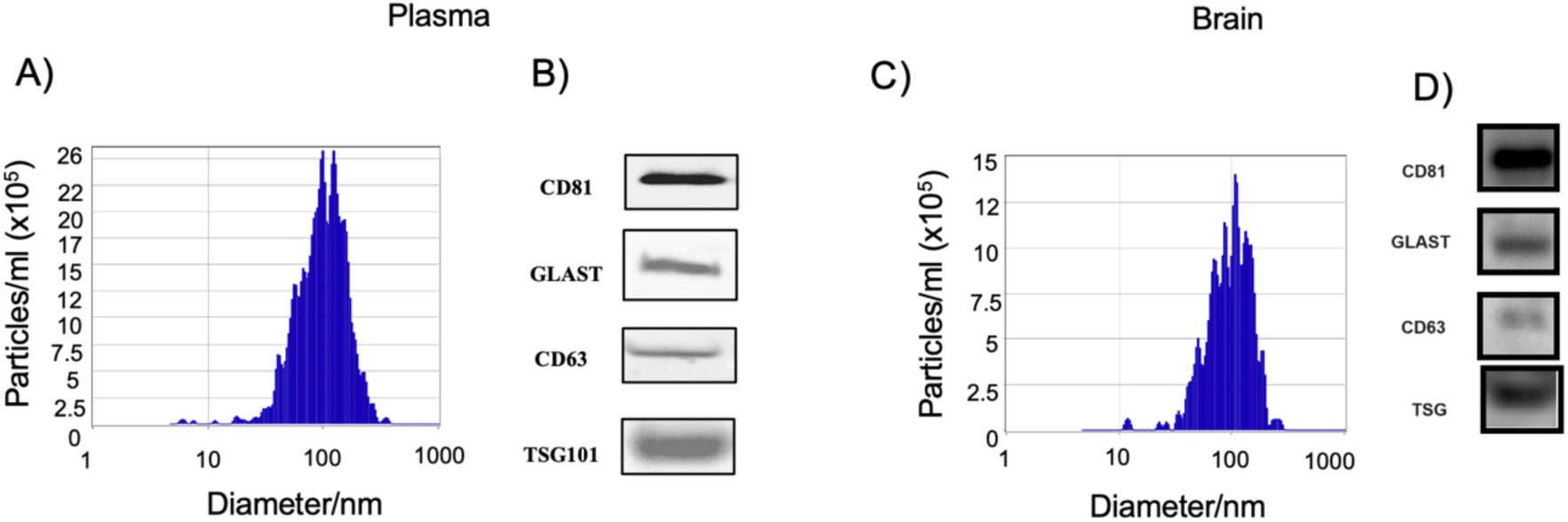
Representative size distribution histogram of exosomes. Total exosomes isolated from plasma and brain from control mouse have been determined by the Zetaview particle analyser (A, C) and by Western-blot of CD81, Glast, CD63 and TSG markers (B, D).

**Supplemental figure 2:**
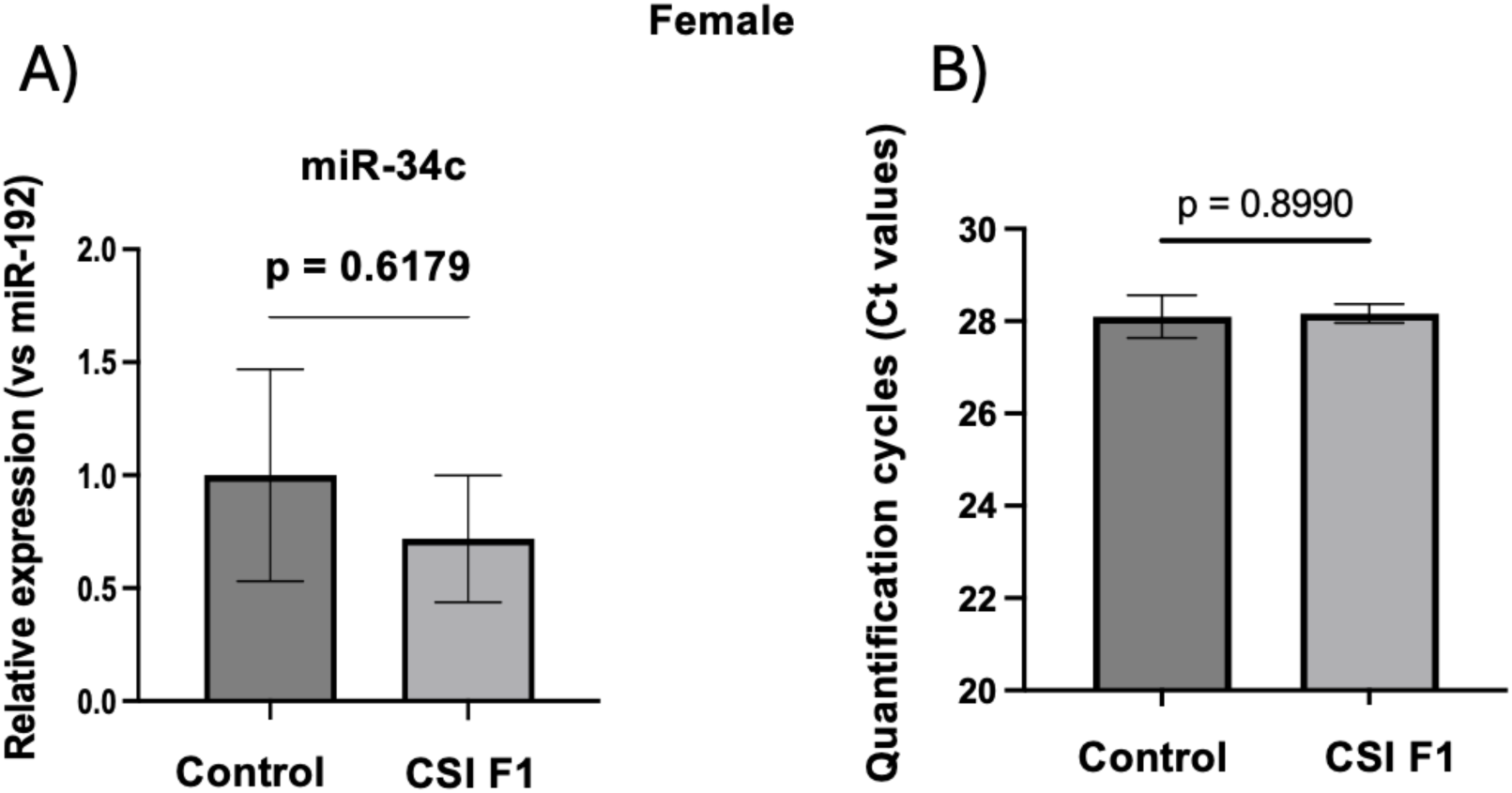
Astrocytes-exosomes miR-34c in F1 CSI females. Astrocytes-exosomes (A-Exos) are isolated form plasma females F1 issued of males control of CSI stressed. The level of miR-34c is unchanged in A-Exos (A). Level of miR-192 (Ct value) in A-Exos from females F1. Data are expressed as mean ± S.E.M; T-test All values are compared to the internal control miR-192 that does not change enough by any perturbation to significantly alter the conclusions and control ratios are set to 1. N=7 for control and N=8 for CSI F1. All values are compared to the internal control miR-192 that does not change enough by any perturbation to significantly alter the conclusions and control ratios are set to 1.

**Supplemental figure 3:**
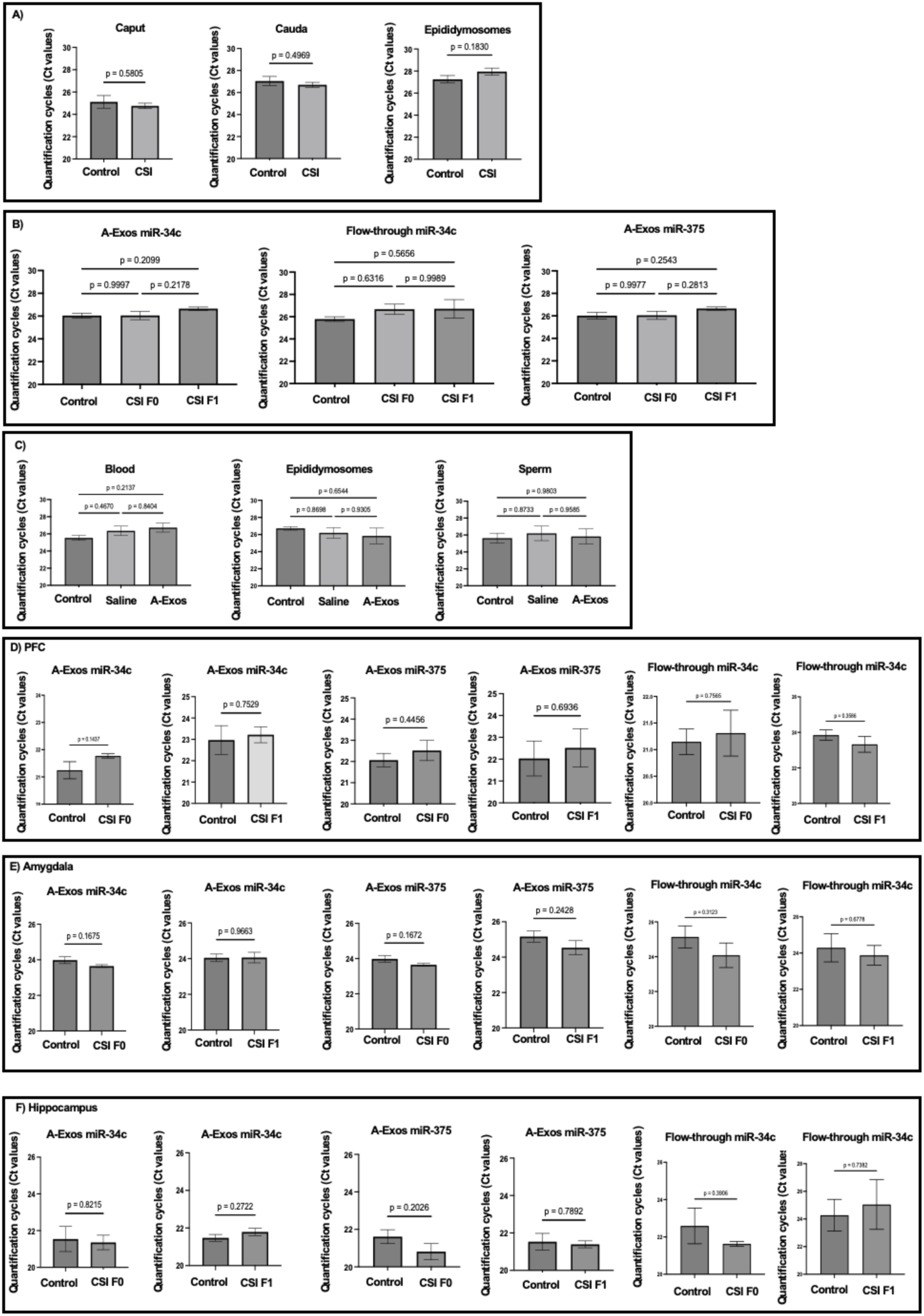
Quantification of cycle of internal normalizer. A) Quantification cycles of miR-192 from figure 1. B) Quantification cycles of miR-192 from figure 2. C) Quantification cycles of miR-192 from figure 3. D) Quantification cycles of miR-192 from figure 4 on PFC. E) Quantification cycles of miR-192 from figure 4 on Amygdala. F) Quantification cycles of miR-192 from figure 4 on Hippocampus. Data are shown as mean of Ct values +/- SEM. One-way Anova with multiple comparisons for 3 groups (figures 2 and 3) and T-test for figures 1 (A) and 4 (D-F). PFC: prefrontal cortex.

## Notes

### Competing Interest Statement

The authors have declared no competing interest.

### Summary of Updates

minor editing of introduction where the end of a sentence was omitted.

